# Detection of regularities in auditory sequences before and at term-age in human neonates

**DOI:** 10.1101/2023.08.15.553237

**Authors:** Marine Panzani, Mahdi Mahmoudzadeh, Fabrice Wallois, Ghislaine Dehaene-Lambertz

## Abstract

During the last trimester of gestation, fetuses and preterm neonates begin to respond to sensory stimulation and to discover the structure of their environment. Yet, neuronal migration is still ongoing. This late migration notably concerns the supra-granular layers neurons, which are believed to play a critical role in encoding predictions and detecting regularities. In order to gain a deeper understanding of how the brain processes and perceives regularities during this stage of development, we conducted a study in which we recorded event-related potentials (ERP) in 31-wGA preterm and full-term neonates exposed to alternating auditory sequences (e.g. “ba ga ba **ga ba**”), when the regularity of these sequences was violated by a repetition (e.g., "ba ga ba **ga ga**"). We compared the ERPs in this case to those obtained when violating a simple repetition pattern (“ga ga ga **ga ga**” vs “ga ga ga **ga ba**”). Our results indicated that both preterm and full-term neonates were able to detect violations of regularity in both types of sequences, indicating that as early as 31 weeks gestational age, human neonates are sensitive to the conditional statistics between successive auditory elements. Full-term neonates showed an early and similar mismatch response (MMR) in the repetition and alternating sequences. In contrast, 31-wGA neonates exhibited a two-component MMR. The first component which was only observed for simple sequences with repetition, corresponded to sensory adaptation. It was followed much later by a deviance-detection component that was observed for both alternation and repetition sequences. This pattern confirms that MMRs detected at the scalp may correspond to a dual cortical process and shows that deviance detection computed by higher-level regions accelerates dramatically with brain maturation during the last weeks of gestation to become indistinguishable from bottom-up sensory adaptation at term.

**Highlights:**

- Starting at 31 wGA, neonates are sensitive to conditional statistics between successive events.
- The MisMatch Response detected at the scalp may correspond to a dual cortical process
- The prediction error signal accelerates during the third trimester of gestation
- It overlaps with the phenomenon of sensory adaptation at term age

## Introduction

Full-term birth in humans occurs after 38 weeks of gestation (wGA). The last trimester of human gestation is marked by ongoing neuronal migration to create the characteristic six-layer cortex. Most neurons lie beneath the cortical plate, creating a thicker compartment clearly visible by MRI (Kostovic & Vasung, 2009) that slowly disappears until a few weeks post-term in the frontal areas (Paredes et al., 2016). However, neural activity is recorded from the first waves of neurons arriving in place and evoked activity to external events is observed from 28 wGA, when thalamic afferents enter the cortical plate. At this age, premature neonates not only respond to sounds but are already able to detect rhythm violations (Edalati et al., 2022), distinguish phonemes (ba vs ga) and voices (male vs female) by exploiting different cortical networks. Notably, phonetic discrimination is supported by an extended peri-sylvian network from the superior temporal region to the inferior frontal gyrus (Mahmoudzadeh et al., 2013), revealing functional bottom-up pathways to higher-level regions. Despite ongoing neuronal migration, which, during the last weeks of gestation, mainly concerns supra-granular layer neurons and inter-neurons, learning begins and at term, newborns recognize their native language (Mehler et al., 1988), their mother’s voice (DeCasper & Fifer, 1980; Beauchemin et al., 2011), and melodies to which they were exposed *in utero* (Granier-Deferre, Bassereau, Ribeiro, Jacquet, & Decasper, 2011). However, little is known about the learning mechanisms of this specific period, during which cerebral networks are incomplete, and on how fetuses and preterm neonates begin to discover the regularities of their environment.

Unlike traditional models that rely primarily on bottom-up responses, current influential models of learning (Friston, 2005; Tenenbaum et al., 2011) postulate more complex circuitry that compares inputs with an internal model and reacts when a prediction error occurs to improve the current model. This conceptual shift can be seen in the proposed models of the mismatch response, an electrophysiological component that is evoked when there is a change in stimulus after a series of identical sounds. A first proposal is based on the notion of sensory adaptation in relation to neuronal refractory periods or to weaker firing of the coding neurons due to a sharpening of their response with repetition (Ulanovsky, Las, & Nelken, 2003). After a change in sound, another set of neurons, intermingled with the first set but sensitive to another sound feature - thus not affected by repetition - is solicited, explaining the recovery of the response amplitude at the scalp. Alternatively, other models explain habituation to repetition and the MMR by predictive coding principles involving more complex computations (Friston, 2005; Winkler, 2007; Wacongne et al., 2011; Wacongne, Changeux, & Dehaene, 2012). For example, Wacongne et al. (2012) proposed a neuronal model in which a neuronal population in layer 2-3 would keep track of the regularities of the incoming stimuli to build an internal model, form relevant predictions, and inhibit activity of prediction error neurons in layer 4. Error prediction activity in this layer would occur when the prediction does not match the thalamic input. Predictions might cascade from more-or-less distant regions, depending on the learned regularity, through top-down cortico-cortical connectivity (Opitz, Rinne, Mecklinger, von Cramon, & Schroger, 2002; Wacongne et al., 2011; Sussman, Chen, Sussman-Fort, & Dinces, 2014). MMRs recorded in adults after an omitted but expected event (Wacongne et al, 2012) or after an outlier stimulus that does not belong to a range of many standard stimuli (Garrido, Sahani, & Dolan, 2013) are arguments favoring predictive models over sensory-adaptation. Using MEG, Todorovic & Delange (2012) successfully dissociated the two phenomena in human adults, reporting repetition suppression due to stimulus repetition around 40-60 ms followed by “expectation suppression" around 100-200 ms, both of which being suspended by an unexpected change. Studies in animals using intra-cortical recordings have confirmed that the two phenomena, local sensory adaptation and deviance detection through top-down predictions, might exist together and that both participate in the MMR recorded at the scalp (Chen, Helmchen, & Lütcke, 2015; Lakatos et al., 2020; Teichert, Jedema, Shen, & Gurnsey, 2021).

All these models, except local sensory-adaptation, are based on a precise and specific micro-circuitry involving, notably, the most external cortical layers and interneurons and also efficient cortico-cortical connections, corresponding to a developmental stage that may not be reached before term age. Therefore, investigating the ERPs to violations of different types of auditory regularities in preterm neonates might provide new insights on the neural circuitry supporting deviance-detection mechanisms, clarifying also the learning possibilities before term.

We investigated here whether preterm (∼31 wGA) and full-term (∼40 wGA) neonates can respond to a violation of regularity in simple sequences of five syllables at two levels of sequence complexity: repetition (e.g. ba ba ba ba ba) and alternation (e.g. ba ga ba ga ba) (figure 1). A MMR is recorded after a change in sequences of repeated syllables already at 30 wGA in preterm neonates, (Leppanen et al., 2004; Mahmoudzadeh, et al., 2017), as well as in fetuses (Draganova et al., 2005) but the regularity used in these studies cannot separate the different types of mismatch models. It is why we proposed here to compare the responses in the repetition case to the alternation case. In a sequence built on the alternation between two sounds (ga ba ga ba), the next stimulus (**ga**) is completely predictable once the pattern is discovered. Thus, if the MMR corresponds to genuine detection of the deviance beyond the recovery from sensory adaptation, a deviant sequence, such as ga ba ga **ba ba**, with an unexpected repetition, should elicit a response similar to the classical MMR recorded when the new sound **ga** is presented after several repetitions of **ba** (e.g. ba ba ba **ba ga**). This is what is observed in adults (Wacongne et al., 2012). In our design, the MMR should thus be similar in terms of latency and topography for both types of blocks, yielding a main effect of deviance without an interaction between condition (standard x deviant trials) and type of block (repetition vs alternation). On the other hand, if the MMR is only dependent on sensory adaptation, it should be recorded only in the repetition blocks. In alternation blocks, because both neuronal populations sensitive to ba and ga have been similarly stimulated before the 5^th^ syllable, no difference between standard and deviant trials should be recorded. A final, but less supported possibility is that EEG recordings are sufficiently sensitive to a local repetition (i.e., sensory adaptation after only one item repetition is measurable) and the difference (deviant-standard) changes the sign, depending on the block, because the final transition […ba ga] is deviant in repetition block but standard in alternation blocks, and the opposite for [… ba ba], standard in repetition blocks but deviant in the alternation blocks. This should result in a significant interaction conditioned by the type of block. With this paradigm, we can thus contrast a bottom-up response only leading to a decrease in amplitude (repetition suppression), dependent on the frequency of the heard items in the close past, from more complex computations that require a longer window of temporal integration and eventually predictive processes to take place.

**Figure 1:**
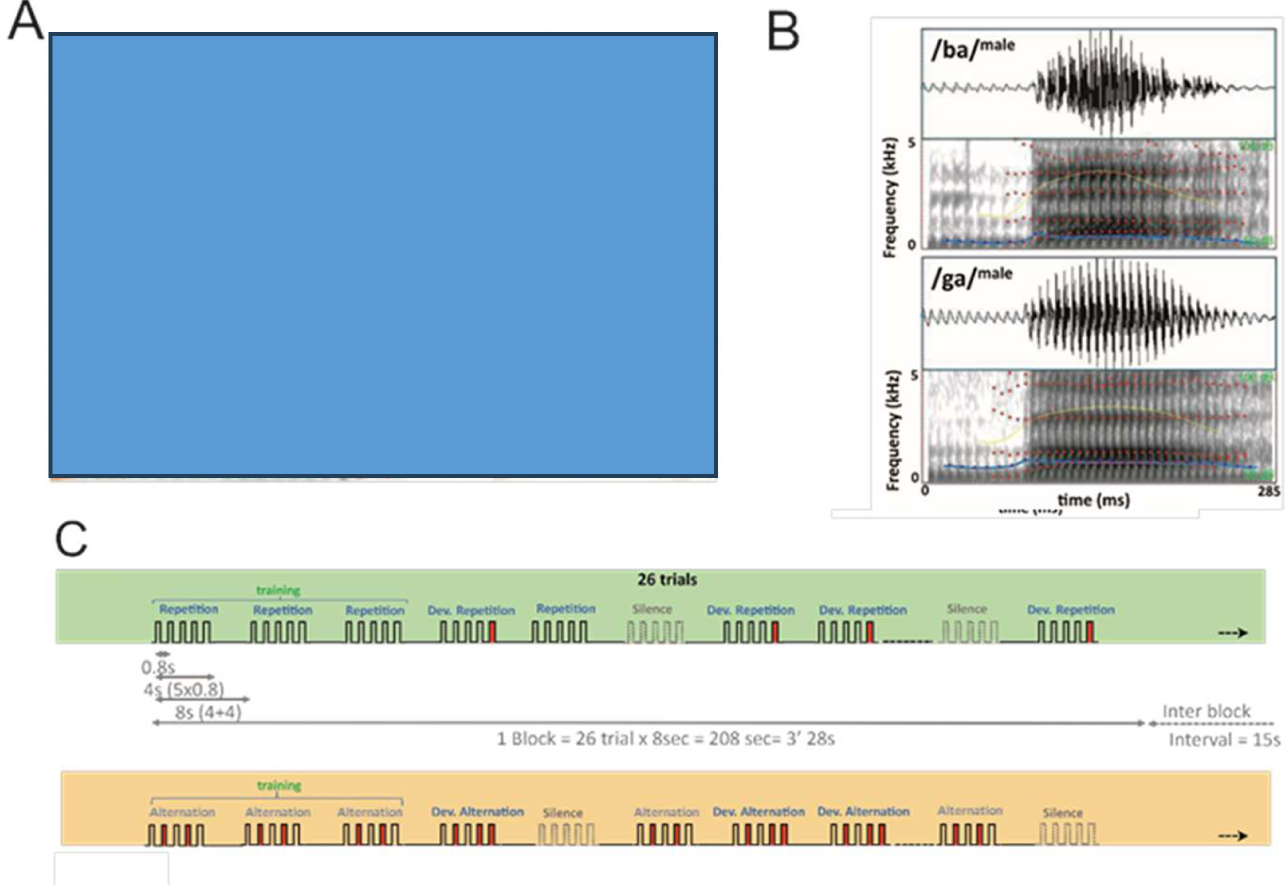
Experimental design. A) A preterm neonate with the high-density net. B) Spectrograms of the two syllables. C) Time-line of the trials in a repetition and an alternation block.

Predictive processes are clearly observed after term birth: At six months post-term, an absent but expected visual stimulus induces an occipital activation visible with near-infrared spectroscopy (Emberson, Richards, & Aslin, 2015). In younger infants, predictive capacities have been studied through violations of repeated patterns. Three-month-old awake infants (Basirat, Dehaene, & Dehaene-Lambertz, 2014), as well as full-term neonates and fetuses after 35 weeks of gestation (Moser et al., 2020), exposed to sequences such as AAAB AAAB AAAB, … showed a mismatch response when an expected change at the end of the sequences is not present. In other words, a strict repetition (AAA), if not expected, evokes a MMR.

From these results, we expected that full-term neonates should learn the two regularities (repetition and alternation) and present similar responses in terms of habituation and change detection in the two types of blocks. At 31 wGA, neuronal migration is still ongoing and the intra-cortical connections immature. Thus, preterm neonates might only react to each sound with the active neuronal population at that time (i.e., those not in the refractory period induced by the previous sound) and thus present a MMR explained by sensory adaptation only in the repetition blocks. On the other hand, they might also react in the same way as full-term infants, thus requiring new anatomical models to explain the brain’s predictive abilities. Indeed, the failure to react to an AAAA sequence in blocks of AAAB sequences, in younger fetuses reported by Moser et al. (2021) might be due to the challenge of fetal MEG and preterm neonates are easier to test under the exact same conditions as full-term infants, offering the possibility to study how learning possibilities progress with brain development and setting-up of the cortical layers.

## Materials and Methods

### 1 Participants

#### Full term neonates

Twenty healthy full-term neonates (10 males, 10 females) were recorded but five neonates were excluded due to the recording being too short (3) or a technical problem (2). For the 15 remaining infants (7 males, 8 females), the mean GA at birth was 39.8 ± 1.2 weeks, mean birth weight: 3365 ± 442 g, mean cranial circumference 34.6 ± 1 cm, Apgar score = 10 at 5 min, and mean recording age: 3.5 ± 1days (see Table 1 for details).

**Table 1:**
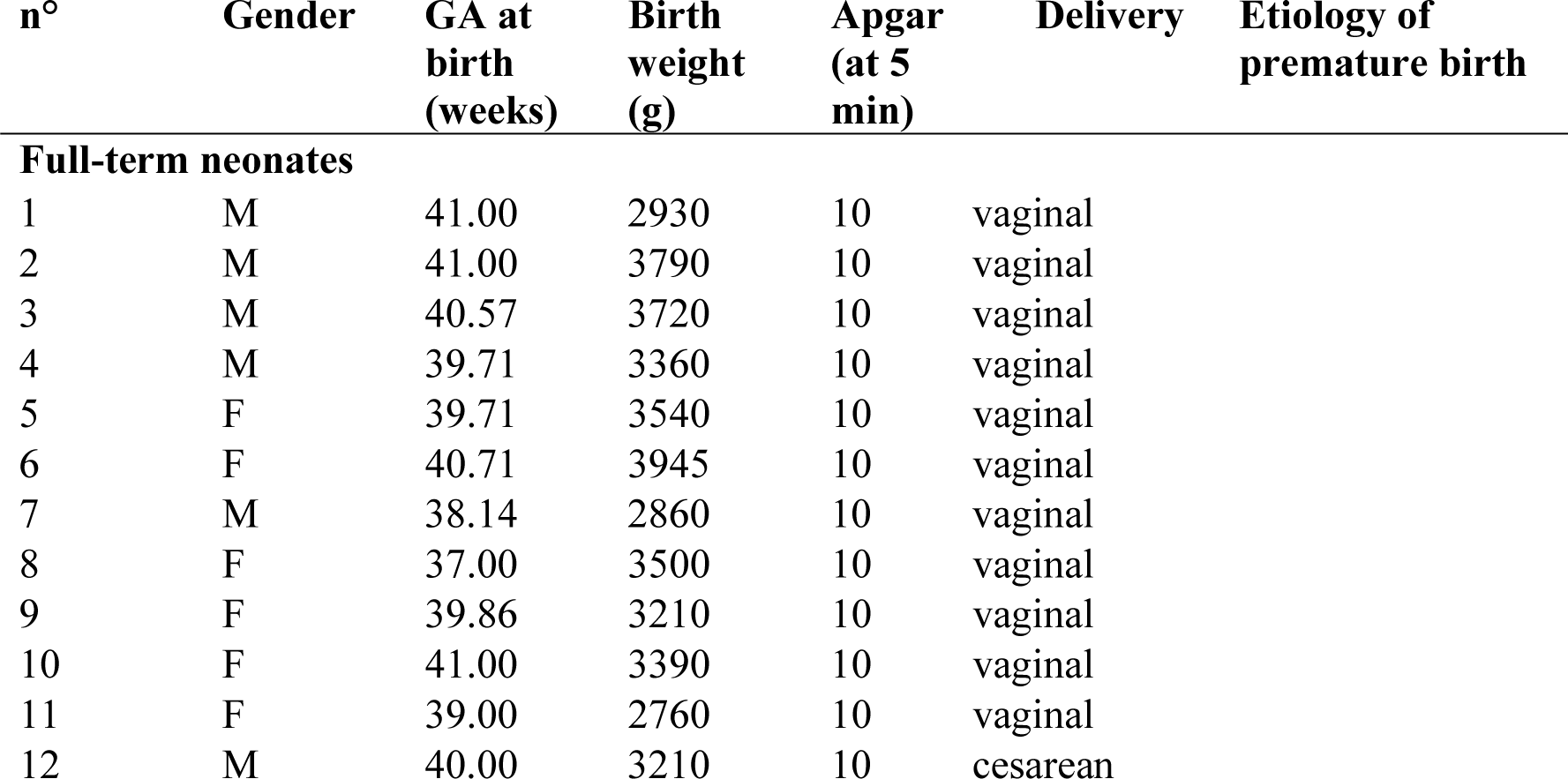
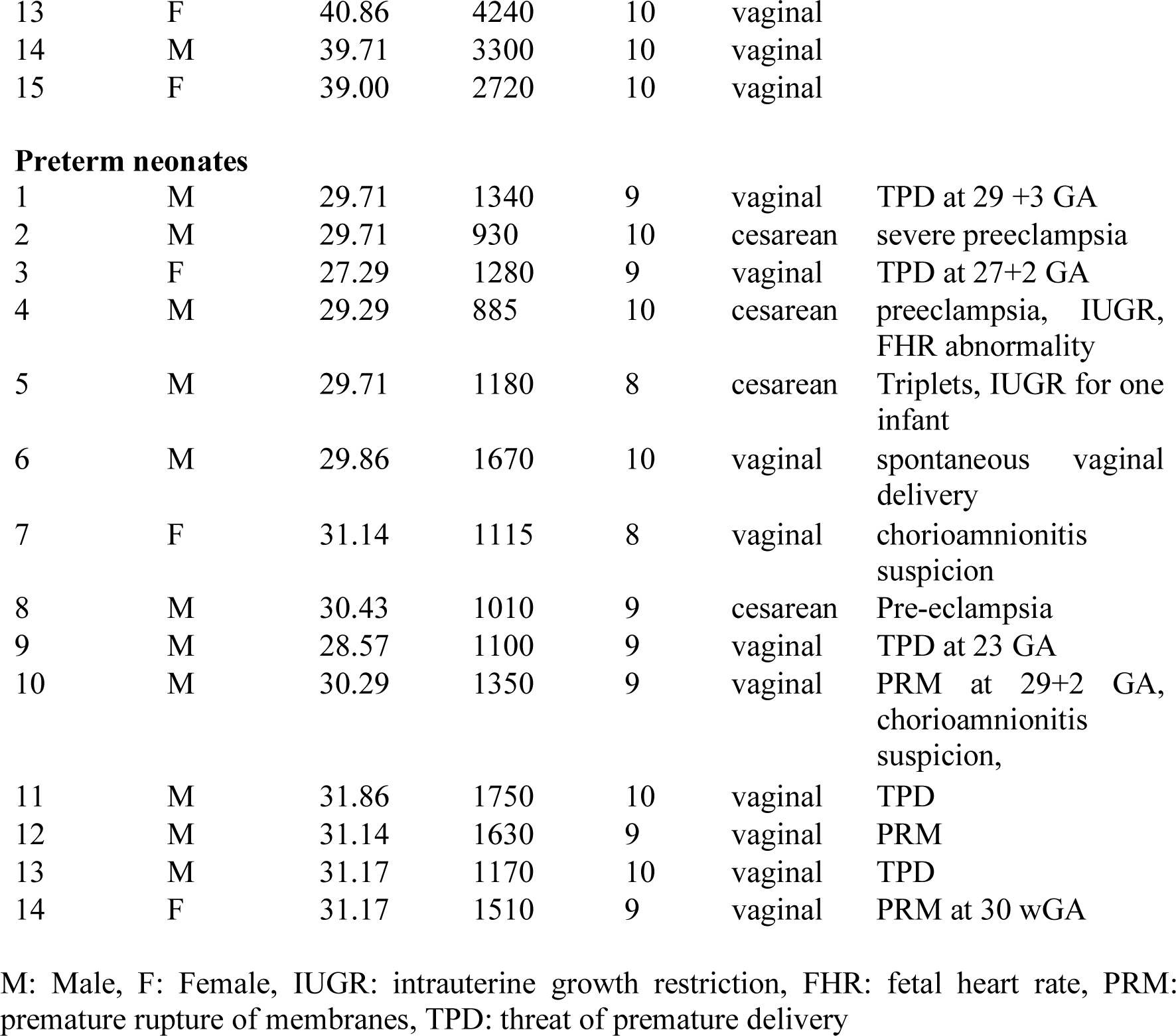
Characteristics of the two groups.

| n° | Gender | GA at birth (weeks) | Birth weight (g) | Apgar (at 5 min) | Delivery | Etiology of premature birth |
| --- | --- | --- | --- | --- | --- | --- |
| **Full-term neonates** |  |  |  |  |  |  |
| 1 | M | 41.00 | 2930 | 10 | vaginal |  |
| 2 | M | 41.00 | 3790 | 10 | vaginal |  |
| 3 | M | 40.57 | 3720 | 10 | vaginal |  |
| 4 | M | 39.71 | 3360 | 10 | vaginal |  |
| 5 | F | 39.71 | 3540 | 10 | vaginal |  |
| 6 | F | 40.71 | 3945 | 10 | vaginal |  |
| 7 | M | 38.14 | 2860 | 10 | vaginal |  |
| 8 | F | 37.00 | 3500 | 10 | vaginal |  |
| 9 | F | 39.86 | 3210 | 10 | vaginal |  |
| 10 | F | 41.00 | 3390 | 10 | vaginal |  |
| 11 | F | 39.00 | 2760 | 10 | vaginal |  |
| 12 | M | 40.00 | 3210 | 10 | cesarean |  |

|  |  |  |  |  |  |
| --- | --- | --- | --- | --- | --- |
| 13 | F | 40.86 | 4240 | 10 | vaginal |
| 14 | M | 39.71 | 3300 | 10 | vaginal |
| 15 | F | 39.00 | 2720 | 10 | vaginal |
| **Preterm neonates** |  |  |  |  |  |  |
| 1 | M | 29.71 | 1340 | 9 | vaginal | TPD at 29 +3 GA |
| 2 | M | 29.71 | 930 | 10 | cesarean | severe preeclampsia |
| 3 | F | 27.29 | 1280 | 9 | vaginal | TPD at 27+2 GA |
| 4 | M | 29.29 | 885 | 10 | cesarean | preeclampsia, IUGR, FHR abnormality |
| 5 | M | 29.71 | 1180 | 8 | cesarean | Triplets, IUGR for one infant |
| 6 | M | 29.86 | 1670 | 10 | vaginal | spontaneous vaginal delivery |
| 7 | F | 31.14 | 1115 | 8 | vaginal | chorioamnionitis suspicion |
| 8 | M | 30.43 | 1010 | 9 | cesarean | Pre-eclampsia |
| 9 | M | 28.57 | 1100 | 9 | vaginal | TPD at 23 GA |
| 10 | M | 30.29 | 1350 | 9 | vaginal | PRM at 29+2 GA, chorioamnionitis suspicion, |
| 11 | M | 31.86 | 1750 | 10 | vaginal | TPD |
| 12 | M | 31.14 | 1630 | 9 | vaginal | PRM |
| 13 | M | 31.17 | 1170 | 10 | vaginal | TPD |
| 14 | F | 31.17 | 1510 | 9 | vaginal | PRM at 30 wGA |

M: Male, F: Female, IUGR: intrauterine growth restriction, FHR: fetal heart rate, PRM: premature rupture of membranes, TPD: threat of premature delivery

#### Preterm neonates

Sixteen preterm neonates (5 males, 12 females; mean GA at birth: 29.8 ± 2 weeks) were tested during the first two weeks of life, before 33 wGA. All had normal clinical neurological assessments, a normal EEG for term age, and a low risk of brain damage. Two neonates were excluded due to technical problems during the recording. For the 14 remaining infants (11 males, 3 females), the mean GA was 30.1 ± 1.2 weeks at birth and 31.4 ± 1.7 weeks at the time of the test, corresponding to 11.7± 6 days of age. The Apgar score was >= 8 at 5 min, mean birth weight: 1280 ± 282 g (see Table 1 for details).

All parents were informed about the study and provided their written informed consent. The study was approved by the local ethics committee (CPP Nord-Ouest II; 2008-A00728-47).

### 2 Stimuli and Experimental Paradigm

The same two syllables produced by a French male speaker as in the study of Mahmoudzadeh et al. (2013) (/ba/ and /ga/) were used (Fig. 1B). They were matched for intonation, intensity, total duration (285 ms), prevoicing, and voiced formant transition duration (40/45 ms). They were presented at a comfortable hearing level (≈70 dB) by series of five syllables (SOA = 800 ms) to form four types of trials (standard and deviant trials in a repetition or alternation context). The first four syllables created the context. The same syllable (/ba/ or /ga/, depending on the block) was repeated four times in the repetition context followed by itself in standard trials (e.g., ba ba ba **ba ba**) or by the other syllable in deviant trials (e.g., ba ba ba **ba ga**), breaking the repetition pattern. In the alternation context, the syllables alternated between ba and ga (/ba/ga/ or /ga/ba/, depending on the block), the last and 5^th^ syllable following the alternation rule in standard trials (e.g., ga ba ga **ba ga**), or repeating the 4^th^ in deviant trials and breaking the pattern (e.g., ga ba ga **ba ba**). The intertrial interval between the auditory sequences was 4 000 ms.

Each infant received every type of trial, blocked according to the trial pattern (alternation or repetition, two blocks of each presented, interleaved, Fig.1C). Each block (duration 208 s) comprised 26 trials and was followed by 15 s of silence before the next block. Each block started with a different syllable (/ba/ or /ga/), thus both syllables were similarly presented at each standard and deviant position. Blocks consistently commenced with three standard trials to establish the block pattern, then 10 standard, 10 deviant, and three silence trials were randomly intermixed. The silent trials, in which a silence of the same duration (4s) replaced the sequence of syllables, were needed because it was initially planned to simultaneously used near-infra-red spectroscopy (NIRS) and EEG to record the brain responses. Infants listened to 16 blocks for a total of 416 trials, comprising 48 silent trials (total duration: 59.5 min).

The experimental protocol was coded in Matlab®. Synchronization between the stimulation computer and the EEG acquisition device was performed via an external trigger. The sound was presented through speakers (RAL ELEVATE3 ®; 80Hz – 20kHz) to full-term neonates and B&O PLAY Beoplay A1 ®; 60Hz-24KHz) to preterms who were tested inside their incubator, with the speaker at the infant’s feet. The stimuli were presented at comfortable hearing level (∼70 dB).

### 3 EEG Recording

Data were recorded using a 128-channel EGI net, adapted to the head circumference, referenced to the vertex, connected to a NetAmps 300 (Electrical Geodesics, Eugene, OR, United States). Before analyses, the reference is transformed to a reference-average and the vertex electrode is reconstructed as the negative mean of all electrodes at each time-point. In the EGI system, there is an “isolated common” on the net, located on the midline between Cz and Pz and “tied to the zero level or common of the isolated amp circuit’s power supply” (Geodesic sensor net technical manual, Philips, p. 101). For four premature infants who had a head circumference smaller than the smallest EGI net (< 28 cm), tissue caps (Malterre® company) in which 64 electrodes (Easy cap ®) were inserted according to the 10-10 system were connected to a SynAmps RT (Compumedics Neuroscan, Charlotte, NC, United States). Data were sampled at 1,000 Hz. The electrode impedance was kept constant, < 30 kΩ. To keep the sponges moist, the cap was covered with a cellophane film affixed using a Surgifix ® net. Each recording started when the neonate was asleep. A 5-min resting-state period was recorded without stimulation before and after the experiment.

### 4 Data processing

#### Artefact rejection and global averaging

Individual EEG data were bandpass filtered at 0.5-30 Hz (6-24dB/oct; forward and zero phase type, respectively) before artefact rejection (artefact scan tool in BESA Research 6.1, Brain Electrical Source Analysis®). Individual channels were rejected when their absolute amplitude exceeded 50µV or when the gradient exceeded 70µV to reject high-amplitude periods of *trace alternant*, hallmark of preterm EEG. A channel was rejected for the entire session if it was rejected on more than 70% of the trials. The entire trial was rejected if more than 15% of the channels were rejected. On a total of 416 trials, we obtained an average of 198 usable trials for full-term infants and 251 usable trials for preterm infants (Table 2).

**Table 2.**
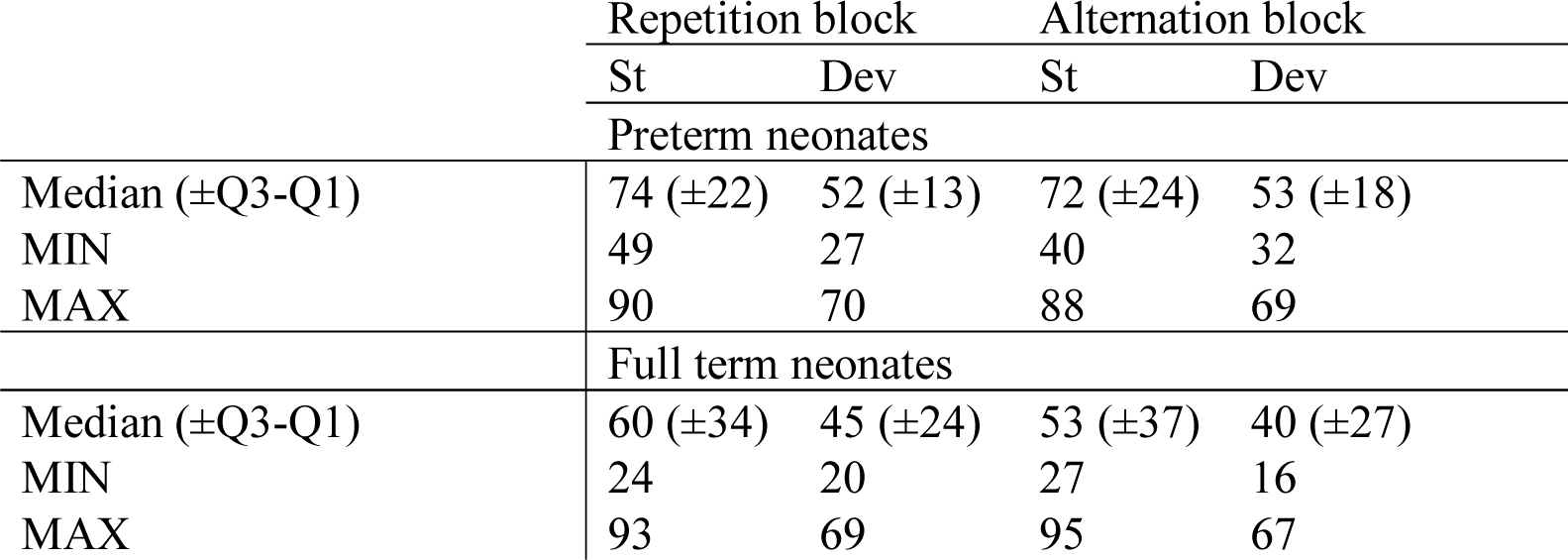
Number of trials for each condition and group.

Due to the different number of electrodes used for the recordings for the premature newborns (64 or 128), data were interpolated to a standard 81 electrodes using the interpolation tool in BESA Research®. Then, individual averaging was band-pass filtered at 1-20 Hz (6-24dB/oct; forward and zero phase type, respectively), down sampled to 256 Hz, segmented in 5.2-s epochs ([−0.2 +5]) time-locked to the first syllable of the trial (S1), reference-averaged and baseline corrected to the 200 ms before the onset of S1. We retained 1800 ms after S5 to study eventual late components which are commonly described for post-term infants (Csibra, Kushnerenko, & Grossmann, 2008; Kouider et al., 2013) and preterm neonates (Mahmoudzadeh et al., 2017) around and after 1000 ms. Because of these long segments, which resulted in slow signal drifts that interfered with the ERP analyses, the high-pass filter was set to 1 Hz, as in many studies at this age (e.g. Mahmoudzadeh et al., 2017; Moser et al., 2020; François et al., 2021).

### 5 Statistical analysis

Statistical analyses were performed in each group separately.

#### Context syllables (S1 to S4)

We first analyzed the responses to the first four syllables. Because they were constant across all trials within a block, we merged the standard and deviant trials. We computed the standardized measurement error (SME) of the grand averages and GFP presented in figure 2 and 4 thanks to a bootstrap procedure as proposed by Luck (2021). Subjects are randomly drawn with replacement the same number of time than the number of subjects in the original data to compute a new grand average and its GFP. This procedure is done 1000 times and the SME corresponds to the standard deviation between these 1000 draws. As in the study of Mahmoudzadeh et al. (2017), we summarized the ERP by the global field power (GFP), which corresponds to the spatial variation across channels at each time-point (Skrandies, 1990; Murray, Brunet, & Michel, 2008; Brunet, Murray, & Michel, 2011). Thus, a GFP maximum corresponds to the time at which the negative and positive poles of an ERP component diverge maximally. Because the ERP is more clearly seen after S1 that followed a longer silence, we identified the GFP maxima (i.e., peaks) corresponding to a change in topography of the grand average after S1 at each age. We averaged the GFP values across a 120-ms time window centered on these peaks in each infant and for each syllable presentation (1 to 4) as we did in Mahmoudzadeh et al (2016). We subjected the values for peak 2 and 3 to two ANOVAs with syllable (S1 S2 S3 S4) and block (repetition vs alternation) as within-subject factors. Post-hoc tests were FDR corrected. Because the repetition effect is maximal between S1 and S2 (Dehaene-Lambertz & Dehaene, 1994; Mahmoudzadeh et al., 2016), we performed a similar ANOVA restricted to the two first syllables to investigate whether the voltage decrease was similar, or not, when there was a change in the syllable (AB in alternation blocks) or when it was repetition (AA in repetition blocks).

**Figure 2:**
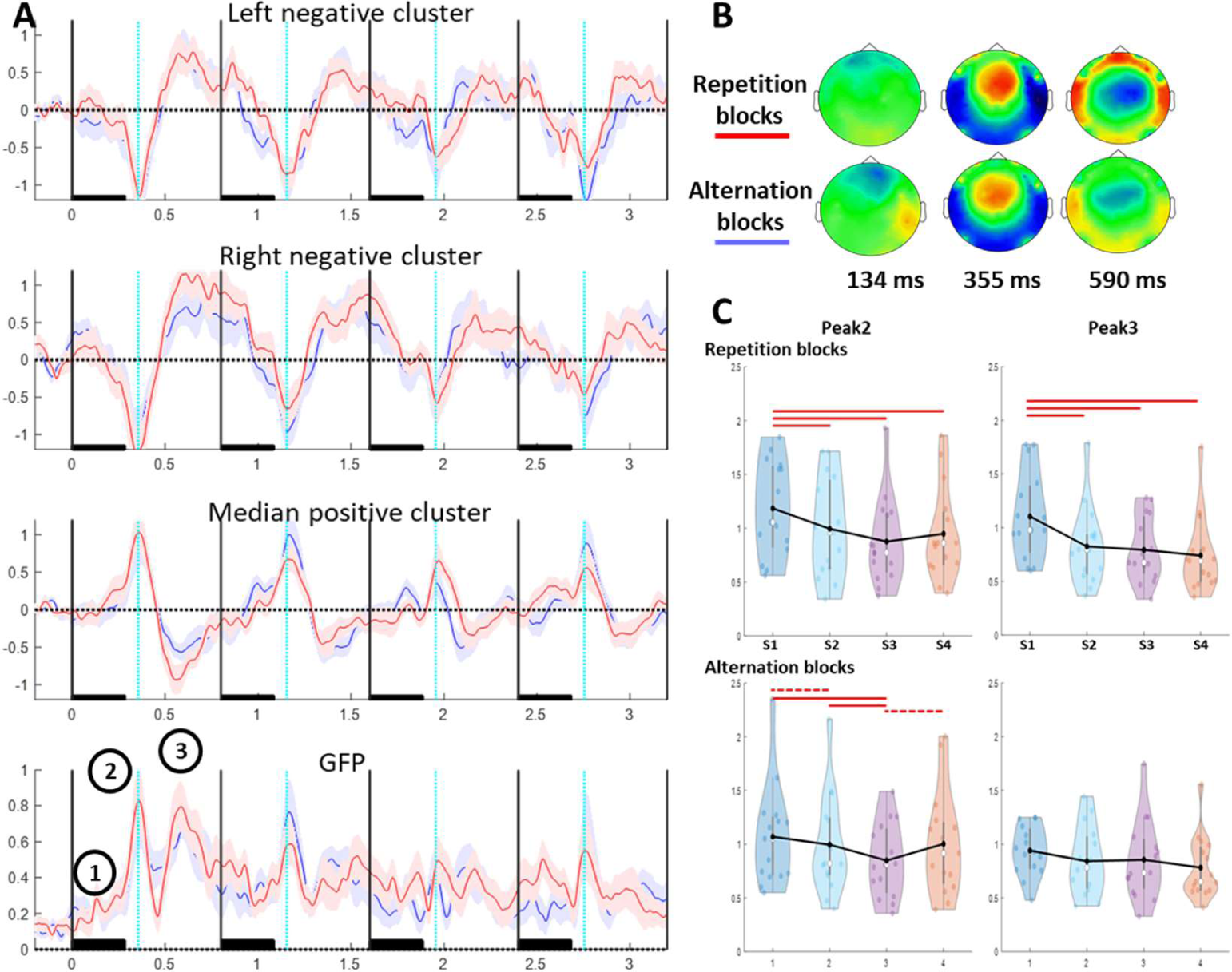
Grand average during the context syllables (S1-S4) in fullterm neonates in Repetition and Alternation blocks. A) the three first rows correspond to the voltage averaged across the channels covering the negative and positive poles of peak 2 and the last row to the mean GFP values. The shaded area around the mean value corresponds to the standard measurement error of the mean, repetition blocks in red and alternation blocks in blue. The vertical black lines indicate the onset of the syllables, for which the duration is indicated by the black rectangle. The cyan dotted line corresponds to the latency of peak 2 (355 ms after each syllable onset). B) Voltage topographies at the peaks of the ERP identified on S1 (circled numbers on GFP plot) for the repetition and alternation blocks. C) Whisker plots superimposed on density-plots of the GFP values around the ERP for peaks 2 and 3 for each of the four syllables (S1 to S4) of the trials. Each point represents a neonate and the white circle the median. The black lines join the mean. The plain red lines indicate the significant comparisons between syllables (p < 0.05) and the dotted lines a trend (0.05 < p < 0.1). The voltage significantly decreased between S1 and the following syllables in repetition blocks. A similar, albeit weaker, pattern was noted for Peak 2 during the alternation blocks.

#### Response to incongruent syllables (S5)

Because of the high number of samples and electrodes, we relied on spatio-temporal cluster-based statistics (Oostenveld, Fries, Maris, & Schoffelen, 2011). This consists of a two-stage analysis, in which t-tests between the studied conditions are first computed for each time-point and electrode. Neighboring time-samples and electrodes that are above a certain threshold (here alpha = 0.05) are grouped into clusters and their t-values added. In the second stage, 1000 permutation tests are performed in which condition labels are randomly shuffled for each participant and the first stage is re-performed for each permutation. We thus obtained the null distribution of the cluster values for the considered comparison. The final probability value corresponds to the number of permutations with t-values that are higher than those obtained in the original data (Maris & Oostenveld, 2007). It should be noted that while this procedure elegantly handles the problem of multiple comparisons in high-density recordings, it cannot provide the exact onset, offset and spatial extent of significant differences between conditions, as the null hypothesis is based on cluster values and does not make any statistical inference on individual samples (Oostenveld et al., 2011; Sassenhagen & Draschkow, 2019).

We considered a long time-window of analysis, 0-1800 ms, to take into account late responses, which are present in infants (Dehaene-Lambertz & Dehaene, 1994; Csibra et al., 2008). In our previous study on infants of this age using only repetition blocks (Mahmoudzadeh et al., 2017), we reported two spatio-temporal clusters between 90-530 ms and 1000-1500 ms post deviance onset. We first performed two comparisons merging all blocks: 1) standard vs deviant trials and 2) local repetition (…AA) vs local change (…AB). Then, we examined the standard vs deviant trials comparison for each type of block and finally considered the Condition (standard vs. deviant) x Block (Repetition vs. Alternation) interaction to capture any difference related to the regularity pattern.

The spatio-temporal cluster-based analysis described above is robust but poorly sensitive. Nor can it provide direct information on the timing of the effect. As we wanted to check whether the MMR might be explained by sensory adaptation, we also carried out an analysis of peak amplitude, which should recover after a syllable change. We therefore considered the same 120ms-time-window centered on Peak 2 and 3 as for habituation analyses and averaged the voltage over the period considered. Paired t-test were computed for each electrode and an FDR correction was applied.

## Results

### 1) Full-term neonates

#### Syllable context (S1 to S4)

Figure 2A presents the grand averages and GFP in full-term neonates in the two types of blocks. The main ERP components were observed at 355 and 590 ms, preceded by a weaker component at 134 ms post-syllable onset. At 355 ms, a central positivity around the vertex was surrounded by a negativity over the temporal and occipital regions, as described by Dehaene-Lambertz and Pena (2001), followed by a reversal in polarity with a central negativity, peaking at 590 ms (Figure 2 B).

As expected from the literature, there was a decrease in amplitude between the first syllable of the block and the following ones visible on the time-series (Figure 2A). To quantify this decrease, we focused on two 120ms-time-windows centered on the GFP maxima at 355 and 590 ms (peaks 2 and 3) (Figure 2C). There was a linear effect of syllables for both peaks (peak 2: F(1,14) = 7.24, p = 0.018; peak 3: F(1,14) = 13.4, p = 0.003), with no effect of the block (F < 1 for both peaks) nor interaction syllable x block (peak 2: F(1,14) = 1.2, p = 0.29; peak 3: F(1,14) = 4.5, p = 0.053).

Pair-wise t-tests between syllables were performed. The GFP values were significantly larger for S1 relative to any other syllables (S2, S3, S4) (all p_fdrcor_ < 0.05) in repetition blocks (AAAAA) for both peaks, without a further decrease between the following syllables. The responses tended to be flatter for alternation blocks (ABABA) compared to repetition blocks. In particular, the ERP to the first syllable was weaker in alternation blocks than in repetition blocks (F(1,14) = 6.65, p = 0.022). Therefore, although a decreasing trend was present between S1 and S2 in alternation blocks (F(1,14) = 3.58, p_fdrcor_ = 0.079), peak 2 was not significantly larger after S1 than after S2 or S4 in these blocks. Anecdotally, syllable 3 evoked a smaller response than all other syllables (S1 vs S3 p_fdrcor_ = 0.043, S2 vs S3 p_fdrcor_ = 0.043, S4 vs S3 p_fdrcor_ = 0.081). The GFP values for peak 3 were flat, with no significant difference between syllables in alternation blocks.

#### Mismatch responses to a change in sequence (S5)

Cluster-based permutation analysis revealed significant differences in the deviant vs standard condition comparison in both types of blocks (Table 3, Figure 3). The difference corresponded to the typical mismatch responses seen at this age, with an anterior positive pole and a reversal of polarity over the posterior regions at the time of the peak of the auditory response (∼350 to 650 ms from the onset of the 5^th^ syllable). No significant cluster was observed in the Condition (standard vs deviant) x Block (alternation vs repetition) interaction. The comparisons within each block are presented in Table 3. The comparison of all trials ending with a repetition (i.e., the 4^th^ and 5^th^ syllables were identical) and all trials ending with a change (i.e., the 4^th^ and 5^th^ syllables were different) showed no significant clusters, regardless of the block.

**Figure 3:**
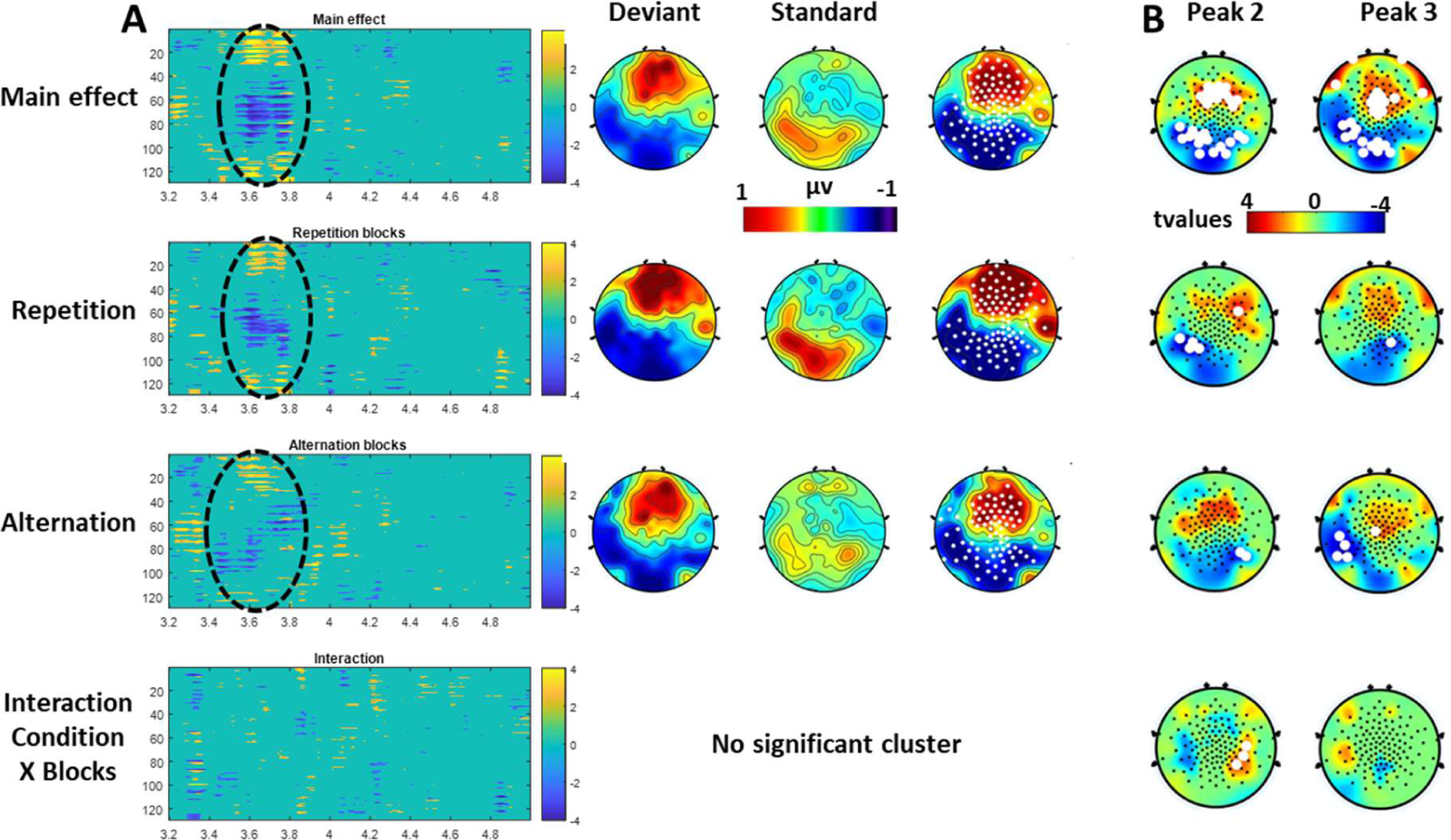
MMRs in full-term neonates. Left column: spatio-temporal clusters of t-test values computed for each electrode and time-sample from the onset of the 5th syllable (3.2 s from the onset of the trial) for the different statistical contrasts. Black circles are manually superimposed over the clusters that are significant relative to the null-distribution obtained over 1,000 permutations (Table 2 for details). On the right, 2D topographies of the mean voltage during the time-window of the significant clusters for the deviant and standard conditions and their difference. Electrodes of the significant clusters are identified by white dots. B) Topographies of t-values from comparison of standard vs deviant trials performed on the voltage averaged across a 120ms-time-window centered on peak 2 (355 ms post S5 onset) and peak 3 (590 ms post S5 onset). White dots represent significant electrodes after FDR correction (p<.05). An early MMR was observed for both types of regularity. Only three electrodes show a significant trials X Blocks interaction for peak 2 due to a different negative pole of the MMR in repetition and alternation blocks.

**Table 3.** MMR clusters in full-term neonates.

| Blocks | Cluster polarity | $p_{\text{cluster-cor}}$ | Time-range* |
| --- | --- | --- | --- |
| All blocks | Positive cluster | $< 0.001$ | 300-671 ms |
| | Negative cluster | $< 0.001$ | 234-632 ms |
| Repetition blocks | Positive cluster | 0.019 | 351-609 ms |
| | Negative cluster | $< 0.001$ | 312-609 ms |
| Alternation blocks | Positive cluster | 0.056 | 230-640 ms |
|  | Negative cluster | 0.039 | 218-741 ms |

\*The time-range indicated correspond to the detection of a significant effect with Cluster-based permutation tests. This method cannot provide the exact onset and duration of an effect

Analyses of S5 peaks confirmed the observation of an early difference in the amplitude of the two peaks for both blocks (Figure 3B). The different topography (a negative pole initially more right lateralized in alternation blocks) induced a significant interaction only during the early time-window (i.e. around peak 2) on three posterior electrodes (p_FDR-corrected_ <0.05).

### 2) Preterm neonates

#### Syllable context (S1 to S4)

In preterm neonates, the main ERP components were identified at 188, 371 and 527 ms post-syllable onset (Figure 4A). The first response at 188 ms consisted of median frontal positivity synchronous with negativity over the posterior electrodes. It was followed by a reversal of the polarity, with two separated anterior negativities synchronous with median posterior negativity peaking at 371 ms, with the anterior negativities then moving medially and the posterior positivity moving laterally, this topography peaking at 527 ms post-syllable onset (Figure 4B). These two last peaks corresponded to the peaks 3 and 4 reported by Mahmoudzadeh et al. (2017) in terms of latency and topography. Concerning the earlier peaks, the match between the present and published data was less obvious, probably due to a difference in the density of recording channels. Peaks with weaker amplitude and/or more focal responses might have been distorted by the small number of recording electrodes and their location in the study of Mahmoudzadeh et al. (2017). Given that all tissues, especially the skull, show higher conductivity in infants, the EEG signal is more focal in infants than later (Grieve, Emerson, Fifer, Isler, & Stark, 2003; Grieve, Emerson, Isler, & Stark, 2004). Thus, the reference average process and the 2D reconstructions of the signal on the scalp are relatively inaccurate if the electrodes do not properly sample the spatial variability in terms of density and overall coverage of the head.

**Figure 4:**
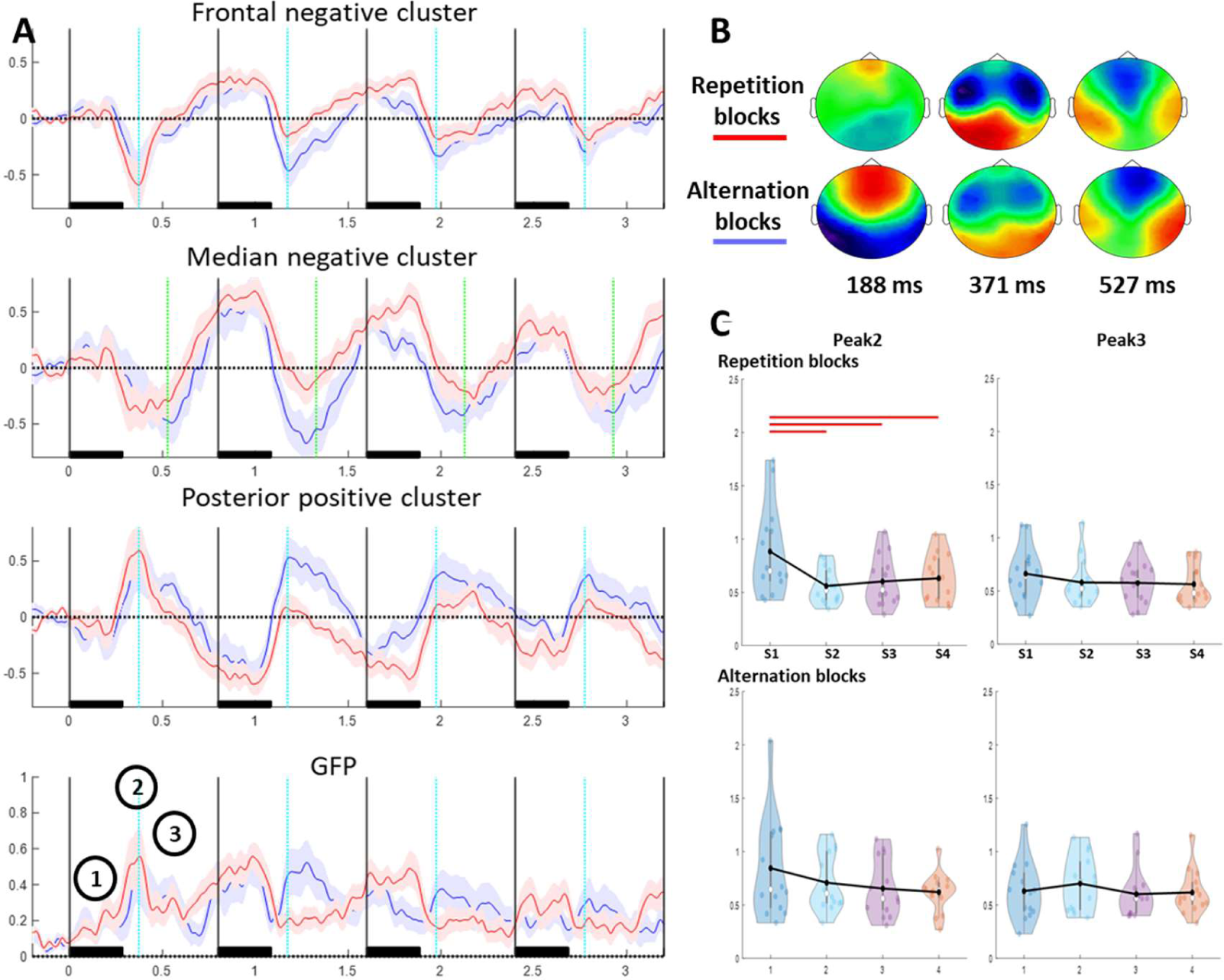
Grand average during the context syllables (S1-S4) in preterm neonates in Repetition and Alternation blocks. A) Rows 1 and 3 correspond to the voltage averaged across the channels covering the negative and positive poles of peak 2, row 2 corresponds to the anterior negative cluster of peak 3, and the last row to the mean GFP values. The shaded area around the mean value corresponds to the standard measurement error of the mean, repetition blocks in red and alternation blocks in blue. The vertical black lines indicate the onset of the syllables, for which the duration is indicated by the black rectangle. The cyan dotted line corresponds to the latency of peak 2 (371 ms after each syllable onset) and the green dotted line in the second plot to peak 3 (527 ms post-onset). B) Voltage topographies at the peaks of the ERP identified on S1 (circled numbers on GFP plot) for the repetition and alternation blocks. C) Whisker plots superimposed on density-plots of the GFP values around the ERP for peaks 2 and 3 for each of the four syllables (S1 to S4) of the trials. Each point represents a neonate and the white circle the median. The black lines join the mean. The plain red lines indicate the significant comparisons between syllables (p < 0.05). The amplitude decrease between S1 and the other syllables is seen only in repetition blocks for peak 2 as it can also been seen in the plots in A.

Similar to the analysis in full-term neonates, we considered two 120ms-time-windows centered on the main GFP maxima at 371 and 527 ms to quantify the habituation (Figure 4C). There was a linear effect of syllables only for the most prominent peak (peak 2: F(1,13) = 5.78, p = 0.032) but not for peak 3 (F(1,13) = 1.64, p > 0.1). There was no effect of block (ps > 0.1) nor a syllable x block interaction (ps > 0.1).

The classical habituation response (i.e., larger response to S1 relative to the subsequent syllables) was only observed for the GFP values of peak 2 in the repetition blocks (all p_fdrcor_ < 0.02). There were no significant differences between syllables for either peak 2 or peak 3 (all p_fdrcor_ > 0.1) for the alternation blocks.

The syllable (S1 vs S2) x block interaction was significant (F(1,13) = 6.8, p = 0.022) due to a block effect for S2 (F(1,13) = 5.85, p = 0.031, corresponding to the expected larger response to S2 in alternation blocks relative to repetition blocks). The decrease in GFP (S1 vs S2) was significant in the repetition blocks (F(1,13) =12.25, p_fdrcor_ = 0.004) but not in the alternation blocks (F(1,13) < 1).

#### Mismatch responses to a change in sequence (S5)

For the main comparison standard vs deviant trials, a late positive cluster was almost significant (p*_cluster-cor_* = 0.055, 921 and 1273 ms). In this latency range, the difference was more pronounced over the right fronto-parietal region (Figure 5). In the analysis restricted to the repetition blocks, two clusters were observed (p*_cluster-cor_* = -0.036: 167 to 607 ms and p*_cluster-cor_* = 0.083: 917-1316 ms). During the first significant latency range, a negative difference between standard and deviant trials developed over the frontal electrodes. Later, the difference became positive and extended progressively clockwise from the frontal region to the temporal region.

**Figure 5:**
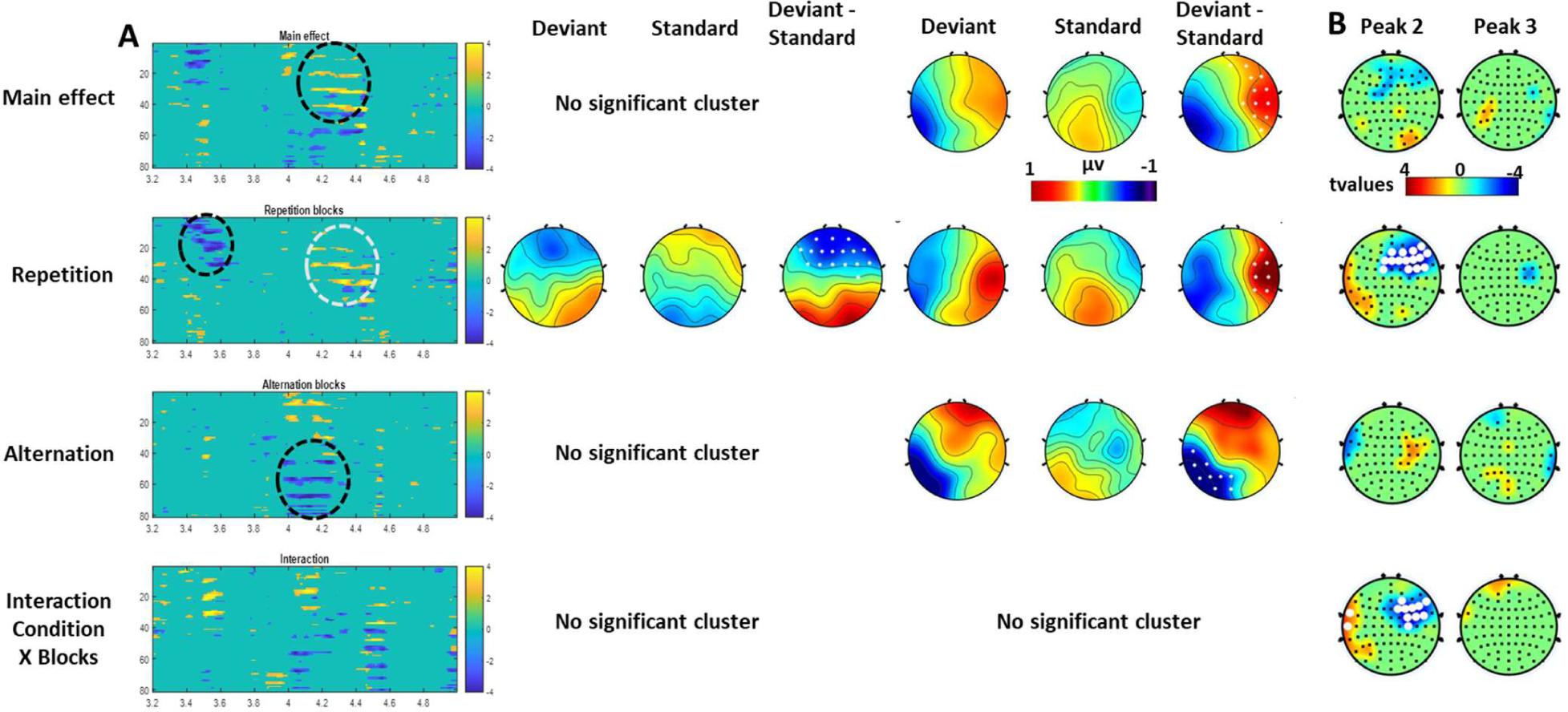
MMRs in pre-term neonates. Left column: spatio-temporal clusters of t-test values computed for each electrode and time-sample from the onset of the 5th syllable for the different statistical contrasts. Black circles are manually superimposed over the clusters that are significant (white circle for a trend) relative to the null-distribution obtained over 1,000 permutations (Table 2 for details). On the right, 2D topographies of the mean voltage during the time-window of the significant clusters for the deviant and standard conditions and their difference. Electrodes of the significant clusters are identified by white dots. B) Topographies of t-values from comparison of standard vs deviant trials performed on the voltage averaged across a 120ms-time-window centered on peak 2 (371 ms post S5 onset) and peak 3 (527 ms post S5 onset). White dots represent significant electrodes after FDR correction (p<.05). There was a significant early difference only for repetition blocks leading to a significant interaction trials X Blocks for peak 2. This first difference is related to sensory adaptation that is possible only in repetition blocks whereas a later cluster corresponded to deviance detection as it is observed for both regularity patterns.

For the alternation blocks, only a late negative cluster (765 to 1089 ms) was significant and corresponded to a posterior negative difference (p*_cluster-cor_* = 0.036) synchronous with an anterior positive difference (p*_cluster-cor_* = 0.17, n.s.). No significant cluster was observed for the interaction Condition (standard vs. deviant) x Block (repetition vs. alternation) interaction.

Comparison of all trials ending with a repetition (i.e., the 4^th^ and 5^th^ syllables were identical) and all trials ending with a change (i.e., the 4^th^ and 5^th^ syllables were different) showed no significant clusters, regardless of the block.

Analyses of S5 peaks confirmed the presence of an early difference solely in peak 2 amplitude between deviant and standard trials in repetition blocks (Figure 5B), leading to a significant interaction with the block type. This suggests that this difference probably arises from a recovery of sensory adaptation that is only visible in response to a syllable change that follows a series of repeated syllables.

## Discussion

We presented two types of auditory regularities to preterm and full-term neonates: either trials consisting of the simple repetition of one syllable (AAA**AA**, e.g., “ba ba ba **ba ba**”) or an alternation between two syllables (BAB**AB**, e.g., “ga ba ga **ba ga**”). We randomly presented violations of the block regularity, obtained by changing the last syllable of the trial (AAA**AB** in repetition blocks, e.g., “ba ba ba **ba ga**”, and BAB**AA** in alternation blocks, e.g., “ga ba ga **ba ba**”). Crucially, the deviant sequence in the alternation blocks consisted of a local repetition (i.e., the 4th and 5th syllables were identical).

For the repetition blocks, we replicated our previous observations at both ages (Dehaene-Lambertz & Dehaene, 1994; Dehaene-Lambertz & Pena, 2001; Mahmouzadeh et al, 2017). First, the repetition of the same sound induced a decrease in the amplitude of the ERP between the first and second sound of the train, with no further decrease between subsequent sounds. Second, a MMR was recorded when a change of sound occurred after a sequence of identical syllables. These results show that, despite the small number of neonates in each group, we had sufficient sensitivity to detect significant effects at each age. They also confirmed in a second study that a subtle phonetic contrast, such as the place of articulation, is already distinguishable at 31 wGA (Mahmoudzadeh et al., 2017). Place of articulation has no invariant acoustic correlates and is easily lost in adverse auditory conditions, such as in a noisy environment, or and in the case of the phonetic impairment observed in dyslexia and aphasia. This ability, replicated in two studies, emphasizes that the human brain is tuned early on, as soon as cortical circuits are being set up, to perceive dimensions useful for language learning.

For the alternation blocks, we observe different results at both ages. Notably, despite slight variations in topography, term neonates exhibited early MMR, regardless of block regularity. By contrast, preterm infants showed an early significant cluster in the repetition blocks, and a late cluster in alternation blocks. Moreover, at the latency of peak 2, only a change of syllable after several repetitions of another syllable produced a modulation of the amplitude.

### Both preterm and full-term neonates learn an alternation regularity

Before discussing the dissimilarities in mismatch responses to regularity violation between preterm and full-term neonates, we would first like to emphasize that at both ages, the neonates reacted to a violation of regularity in alternation blocks, i.e., to a violation that consists of a repetition. This response reveals a genuine deviance detection phenomenon that cannot be reduced to the recovery of sensory adaptation. Indeed, in both the standard and deviant trials of alternation blocks, “ba” and “ga” were equally repeated before the 5^th^ syllable. The equal frequency of the two syllables was also constant throughout the blocks, as the standard (e.g. [ga ba ga ba ga] = 2 ba 3 ga) and deviant trials (e.g. [ga ba ga ba ba] = 2 ga 3 ba) were equally distributed, except for the first three trials of the blocks. Thus, refractory period, neurotransmitter exhaustion in immature neurons or any other bottom-up and neuronal mechanisms that might be affected by the close repetition of the same stimulus cannot explain the difference between standard and deviant trials in the alternation blocks. It was also not due to the single repetition between the 4^th^ and 5^th^ syllable in deviant trials compared to the syllable change at the same position in standard trials because the difference (Deviant – Standard) would have changed of sign relative to repetition blocks. It was not the case neither in the full-term nor the preterm neonates and the comparison …AB vs …AA that directly tested this local repetition effect, did not revealed any significant cluster. Therefore, sensory adaptation is not a mechanism that can explain the detection of a violation of regularity in alternation blocks. This result is in agreement with those obtained by Moser et al (2020, 2021) showing that fetuses above 35 wGA and full-term neonates also reacted to a repetition of four identical tones (AAAA) that broke the regularity of the presentation of AAAB sequences, i.e. sequences of three repeated tones followed by the other tone. What other mechanisms, beyond sensory adaptation, might explained these results?

Sequence regularities can be described, and thus processed, at different levels. For example, alternation trials (ba ga ba ga ba) can be encoded as an abstract rule of alternation between any two syllables (if A then any B), or as transition probabilities between ba and ga (ba predicts ga and ga predicts ba), or even as a long “word” (bagabagaba) held in memory. This last possibility of a memorized word cannot explain our results because the two possible words (e.g. bagabagaba and bagabagaga), corresponding to the standard and deviant trials were presented the same number of times (except for the first three trials of the block) and thus should be similarly familiar to the infants. In the other two cases, a MMR for the deviant trials is explained either because the neonates expected a change that did not occur (alternation rule) or because they reacted to the unexpected transition ba-ba or ga-ga never heard before (statistical learning).

Regarding statistical learning, several studies have indeed shown that neonates, at least at term age, are sensitive to conditional statistics between syllables. For example, they use transition probabilities between syllables to discover words in an artificial stream composed of four randomly concatenated tri-syllabic non-words (Teinonen, Fellman, Naatanen, Alku, & Huotilainen, 2009; Kudo, Nonaka, Mizuno, Mizuno, & Okanoya, 2011; Fló, Benjamin, Palu, & Dehaene-Lambertz, 2021; see also Benjamin et al., 2021 for quadrisyllabic words). Because any of the three other words can follow a given word, there is a drop in transition probability between syllables at the end of the word, from 1 within a word to 0.33 between words. Sleeping full-term neonates are sensitive to this drop. The Moser et al’s studies (2021, 2022) rather favor a rule learning (the last syllable of the sequence should change) but the results were negative under 35 wGA. Note, however, that an alternation rule (first-order regularity) is easier to discover than the second-order regularity used by Moser et al (2021, 2022) because it requires a shorter temporal integration of the auditory input, within the sequence here vs. over multiple trials spanning several tens of seconds in Moser et al’s paradigm. The present experimental paradigm cannot disentangle whether neonates were only reacting to the local transition probabilities or were also able to access a more abstract rule of alternation. This question requires further study. However, the recording of a MMR in alternation blocks in the preterm group reveals that from 31wGA onward, neonates not only react to the present stimulus but are integrating it with the previous auditory history despite incomplete cortical layer formation.

Moser et al. (2020) reported another learning limitation in their study: neonates reacted to the deviant-pattern trials only when they were in an active state, shown by a higher variability in heart-rate during this state. However, another study on associative learning reported better learning during quiet sleep than active sleep. After neonates learned that an air-puff to the eye followed a tone, there was an increase in eye-lid movements following the tone alone, mostly during quiet sleep periods (Tarullo et al., 2016). In the present study, the neonates were tested while asleep but we did not monitor the sleep-stages, as was the case in the studies investigating statistical learning mentioned above. During the last trimester of gestation, active sleep is the first stage to appear, with quiet sleep progressively developing from 28 to 30 wGA, followed by active wakefulness (31 wGA) and then quiet wakefulness (after 35 wGA). The duration of indeterminate sleep stages is still long at 31 wGA (Bourel-Ponchel, Hasaerts, Challamel, & Lamblin, 2021). In full-term infants, sleep cycles typically last 30 to 70 min and consist of 50 to 60% active sleep, ∼40% quiet sleep, and a small amount of indeterminate stage, with frequent transitions between states (Scher, 2008; Bourel-Ponchel et al., 2021). The cycle often starts in active sleep and micro-arousal periods occur within and between sleep states. Thus, the ∼60 min of testing probably included a mixture of different stages at different moments and with different durations between neonates. If anything, the lack of sleep stage identification may have attenuated the MMR. How sleep modulates statistical learning is yet to be investigated, notably in full-term neonates, as well as whether the duration of the integration window for uncovering temporal regularities (over a longer duration in the paradigm of Moser et al. than in the case of alternating syllables) is modulated by the sleep stage.

### The mismatch response: a dual process?

As presented in the introduction, MMRs occurring after a deviant sound within a sequence of repeated sounds have been modeled either as the recovery of sensory adaptation or an error signal between the incoming input and an internal model of the regularity pattern participants are exposed to. Comparing MMRs in repetition and alternation blocks is a way to disentangle these two explanations. In preterm neonates, an early MMR was recorded only for the repetition blocks. This early response replicates the early MMR reported by Mahmouzadeh et al. (2017) in terms of topography (anterior negativity synchronous with posterior positivity) and latency at the same age in an experimental paradigm with only repetition blocks. It also occurred at the latency of peak 2 and with a similar topography (i.e., anterior negativity synchronous with posterior positivity). This early response can be explained by sensory adaptation, with a recovery in the amplitude of peak 2 for the deviant syllable compared to the weaker response to the repeated standard syllable. Sensory adaptation cannot occur for the alternation blocks. There was not even a trend for a MMR during the same early time-window for the alternation blocks. Moreover, when specifically looking at the amplitude modulation of peak 2, a significant Trial x Block interaction was observed with a topography quite similar to the mismatch effect in repetition blocks (Figure 5B). The interpretation of this early difference as a recovery of sensory adaptation following a change of syllable is congruent with the significant decrease in voltage between S1 and S2 in the repetition blocks but not in the alternation blocks in preterm neonates, confirmed by the significant block effect observed for S2.

Later, a second MMR, approximately 1 s post syllable onset, was observed in both type of blocks, with a slightly different topography. The dipole axis was more highly shifted along the left-right axis for the repetition blocks than for alternation blocks. This late response corresponded to the detection of deviance, i.e., an unusual event relative to the stored past.

We thus observed the two phenomena in preterm neonates commonly discussed to explain MMRs, i.e., the release of sensory adaptation (seen in the repetition blocks) and deviance detection (seen in the alternation blocks), generally explained through top-down predictions (Todorovic et al, 2012). Several authors have proposed that instead of the two phenomena being mutually exclusive, both may participate in the MMRs recorded at the scalp because they can be separated in different circumstances (Chen et al., 2015; Lakatos et al., 2020; Teichert et al., 2021). First, certain areas in the auditory cortex only show one or the other effect (Farley, Quirk, Doherty, & Christian, 2010; Fishman & Steinschneider, 2012). Second, in terms of latency, the release of sensory-adaptation after a deviant sound is faster than the deviance detection response (Todorovic et al, 2012; Maheu et al, 2019) as here in preterm neonates. Third, the two mechanisms are differentially sensitive to the inter-stimulus interval (ISI); deviance detection is still present for a long ISI, whereas sensory adaptation decreases with longer ISI (Woods & Elmasian, 1986). Finally, sensory adaptation is released when the new sound is outside the receptive field, whereas deviance detection is more tolerant of small variations and requires larger changes and more repetition to be observed (Lakatos et al., 2020; Teichert et al., 2021).

By contrast, full-term neonates displayed an early MMN, close in amplitude and latency for both the alternation and repetition blocks (Figure 3B). The acceleration of deviance detection probably masks the sensory adaptation phenomenon, which might only weakly contribute to the MMR at term, as it does in adults (Todorovic & de Lange, 2012). The improved integration of the present event into the auditory context is also revealed by the significant decrease in voltage amplitude for both types of blocks in this group (Figure 2). This decrease suggests that at term age, there is a certain level of prediction of what the next syllable will be. It is, nevertheless, paradoxical to note that the difference between blocks in full-term neonates was at S1, which evoked a weaker response in alternation than in repetition blocks. In repetition blocks, S1 represented 90% of the syllables of the block, the other syllable being presented only at the end of the deviant trials, whereas in alternation blocks, the two syllables were equally presented. If the ERP voltage was only modulated by syllable frequency, we might have expected a weaker response for S1 for the repetition blocks. By contrast, the observed stronger response is consistent with a predictive framework, in which highly expected stimuli are primed. The modulation of the S1 ERP by the block regularity, the decrease in ERP amplitude despite the alternating syllables, and, finally, the rapid MMR in both types of blocks reveal a modulation of ERPs by the auditory context in full-term neonates, suggesting efficient and rapid top-down influence on the response to the incoming sensory input.

### Neural changes during the last trimester of gestation

The differences between the two groups is a testimony of the rapid maturation of the cerebral architecture during the last trimester of gestation, but the deviance detection in alternation blocks in preterm infants also underlines that some cortical computations are already allowed before the cortical plate is fully organized. During this period, the microstructures of the cortical columns evolve rapidly in an inside-out manner (Rakic, 1974; Ohtaka-Maruyama., 2015), in which the first neurons to be in place are those of the deep layers. At approximately 28 wGA, most neurons, in particular pyramidal neurons, have migrated towards their final destination (Kostovic, 2020) and thalamic afferents invade the cortical plate after a waiting period in the sub-plate (Kostovic et al., 2019). Although the role of the subplate in modulating neuronal migration is now well understood (Ohtaka-Maruyama et al., 2018; Kostovic et al., 2014), how it contributes to the ERP still requires further exploration (Kostović, 2020). With respect to this issue, the change in polarity of auditory ERPs with maturation has long been known (Vaughan, 1975; Kurtzberg, Hilpert, Kreuzer, & Vaughan, 1984). The reversal of polarity of peak 2 between preterm and full-term infants observed here may correspond to a shift from deep sources to more superficial sources, corresponding to the double circuitry involving the cortical plate and the subplate during the pre-term period followed by middle and upper layer circuitry in the cortical plate later on. Indeed, the subplate, which is particularly prominent in humans, provides a transient circuitry to reinforce the setting-up of the cortical circuitry until the early post-term period. Kostovic (2020) proposed that the subplate has a major role in large-scale interactions between associative areas within and between hemispheres due to the multiple contacts between subplate neurons, interstitial neurons, neurons undergoing migration, and multiple axonal branches of the growing pathways that constitute a complex nexus. This nexus provides additional processing routes that compensate for the weakness of pre-term cortico-cortical connectivity. It could explain the observation in resting-state MRI from 28 wGA onwards of almost all functional networks described later (Fransson et al., 2007; Doria et al., 2010; Fransson, Aden, Blennow, & Lagercrantz, 2011) and probably the extended activated network observed in NIRS when syllables are presented to 28 wGA preterm neonates (Mahmoudzadeh et al., 2013). At term, the cerebral architecture is close to that of later ages, even if remnants of the subplate are still present until the end of the first year of life. Programmed cell death of Cajal-Retzius cells and subplate neurons contribute to this remodeling of the initial circuitry (Molnar, Luhmann, & Kanold, 2020). Furthermore, bottom-up connectivity is pruned away before term, whereas top-down connections are diffuse and pruned only after term (Kennedy, Douglas, Knoblauch, & Dehay, 2007). Thus, although larger responses are recorded in the supra-granular layers in the case of deviance detection in adults (Teichert et al., 2021), the deviance detection response observed in 31-wGA premature neonates beyond sensory adaptation raises new questions about the neural circuitry and, notably, the role of the subplate nexus.

### Limitations of the study

Before to conclude, we would like to point to some limitations of the present study. First the number of neonates per group was small. However, it is worth noting that we successfully replicated the ERP and MMR in terms of latencies and topographies from our previous studies in preterms (Mahmoudzadeh et al., 2016) in which trials AAAA were contrasted with AAAB trials, despite a change of the number of tones in the sequence (4 vs 5 syllables). Additionally, we also replicated previous results in full-term neonates (Dehaene-Lambertz & Pena, 2001). These successful replications supports the reliability of our findings despite the modest sample size. Second, because of the rapid changes in brain size in preterm neonates, we used two different nets of 64 and 128 electrodes, which we converted to a similar template of 81 electrodes by interpolation. This interpolation may have introduced small distortions in the preterms’ voltage topographies. Third, the causes of prematurity are variable and might differently affect the brain even though we excluded the most common and severe conditions such as intraventricular hemorrhage (IVH) and neonates with low APGAR at risk for anoxic injury. Although we cannot completely eliminate brain injury not visible on ultrasound, nor the positive/negative role of the auditory environment on auditory development, care in neonatal units has improved to better manage premature neonates. If we cannot exclude that the latencies reported here may have been affected by deleterious events, it is important to consider that we report here positive evidence of subtle linguistic discrimination on a place of articulation contrast and deviance detection in the case of a sequence regularity based on alternation. These results highlight the early computations an immature brain can perform and thus call for more systematic investigation of the neonatal cognition to be undertaken to improve care.

## Conclusion

Contrary to rodents, in which sensory development occurs after birth, when neuronal migration has finished, sensory responses in primates, notably humans, are superimposed with the neuronal migration stage. Responses to external events are observed not only in primary sensory cortices but also in higher-level regions (Mahmoudzadeh et al., 2013). This situation raises two questions. The first concerns how such external stimulation regulates the last stages of migration and the second, how such an immature network can learn. Here, we have shown that from 31 wGA onwards, neonates are sensitive to an alternation pattern and, thus, at least to the transition probabilities between successive events. Deviance detection is a slow process that is clearly distinct from the bottom-up response eliciting a sensory adaptation phenomenon. During the last two months of gestation, there is a considerable improvement in efficiency, as at term, top-down (deviance detection) processes are predominant in driving the MMR. Investigating how immature humans detect different types of pattern regularities might thus be a powerful tool to reveal the computational properties of the developing connectivity.

## Acknowledgments

This research was supported by a grant to GDL from the European Research Council (ERC) under the European Union’s Horizon 2020 research and innovation program (grant agreement No. 695710), by the French Ministry of Higher Education and Research and by the Regional Council of Picardy (Hemisphere Nord Domi).

## Notes

### Competing Interest Statement

The authors have declared no competing interest.

